# COVID-19 lockdown reveals fish density may be much higher in marine reserves

**DOI:** 10.1101/2022.05.17.492376

**Authors:** Manuel Olán-González, Héctor Reyes-Bonilla, Isabel Montserrat Arreola-Alarcon, Regina Valdovinos Uribe, Damien Olivier

## Abstract

Marine reserves generally allow ecotourism to offer an alternative income to fishing. However, we need to assess its impact on wildlife to make this activity sustainable. The COVID-19 lockdown provided a unique opportunity to evaluate wildlife diversity in the absence of human activity. In a Mexican reserve, we monitored fish assemblages before, during, and just after the lockdown. We show that ecotourism activities alter the behavior of fishes by finding a 2.5-fold density rise during the lockdown. We suggest that the noise pollution generated by the numerous recreational vessels is a significant factor of perturbation. In the absence of noise pollution, some fishes may be bolder (less hidden) and others can come back to the reserve from usually quieter areas (e.g., deeper waters). Our results represent a great worldwide incentive to improve the health of marine reserves by establishing concrete measures in managing plans to mitigate noise pollution.

**Open Research statement:** All data and code necessary to reproduce the results of the paper are enclosed in the submission for review purposes, and will be published on Zenodo following the acceptance of the paper.

## Introduction

During 2020, most countries established social distancing and confinement policies to control the spread of the COVID-19 pandemic. These measures caused changes in human mobility, economic activities, and impacted wildlife and the environment on a global scale (Rutz *et al*. 2020; Montgomery *et al*. 2021). Behavioral and environmental changes were documented around the world, e.g., the presence of wild animals in urban areas (Silva-Rodríguez *et al*. 2020), a reduction in air and water pollution (Zambrano-Monserrate *et al*. 2020), decreases in litter deposited on beaches due to the absence of tourists (Soto *et al*. 2021), and a reduction in noise levels due to the decline in marine traffic (Thomson and Barclay 2020). Therefore, the COVID-19 pandemic created a “Global Human Confinement Experiment” (Bates *et al*. 2020) that offered a unique opportunity to test some effects of human presence and activity on the environment and wildlife.

With a market size valued at $181.1 billion in 2019 and expected to reach $333.8 billion by 2027 (Himanshu and Deshmukh 2021), ecotourism is a growing economic sector attracting more and more visitors to natural reserves. Even though ecotourism provides an economical alternative to resource extraction and is essential to sustaining wildlife conservation (Sala and Giakoumi 2018), human activities and infrastructures associated with this industry can negatively affect the environment. Considering the growth of ecotourism activities, we must evaluate its impact on wildlife to keep it sustainable. However, disentangling the specific effect of the presence of people from other disturbance sources (natural or anthropogenic) is challenging to achieve. Indeed, the closure of natural reserves, which would allow diversity to be assessed in the absence of human activity, even temporarily, is generally not economically viable for the local communities whose livelihood depends on them.

In this study, we used the case of the Cabo Pulmo National Park (“Cabo Pulmo”; 23°26’ N, 109°25’ W, Mexico) to evaluate the effect of ecotourism on the diversity of ichthyofauna. Cabo Pulmo is a 71 km^2^ no-take marine reserve (where any resource extraction is prohibited) and only ecotourism activities have been permitted for more than 20 years (e.g., scuba diving, snorkeling, kayaking). Cabo Pulmo is considered a model well-managed marine reserve because of the local population and government’s strong commitment to its conservation (Rife *et al*. 2013). The management of the reserve has resulted in a 400% increase in reef fish biomass over a decade of protection (Aburto-Oropeza *et al*. 2011). As a result, the improved status of the reef ecosystem and its associated environmental services has attracted people eager to experience swimming in a highly diverse reef where mega-fauna species, such as large fish, sharks, sea lions, and cetaceans, are common (Reyes-Bonilla *et al*. 2014; Cisneros-Montemayor *et al*. 2020). Each month, hundreds of boats carrying thousands of visitors entered the reserve to practice scuba-diving and snorkeling activities (Figure 1a, b).

**Figure 1.**
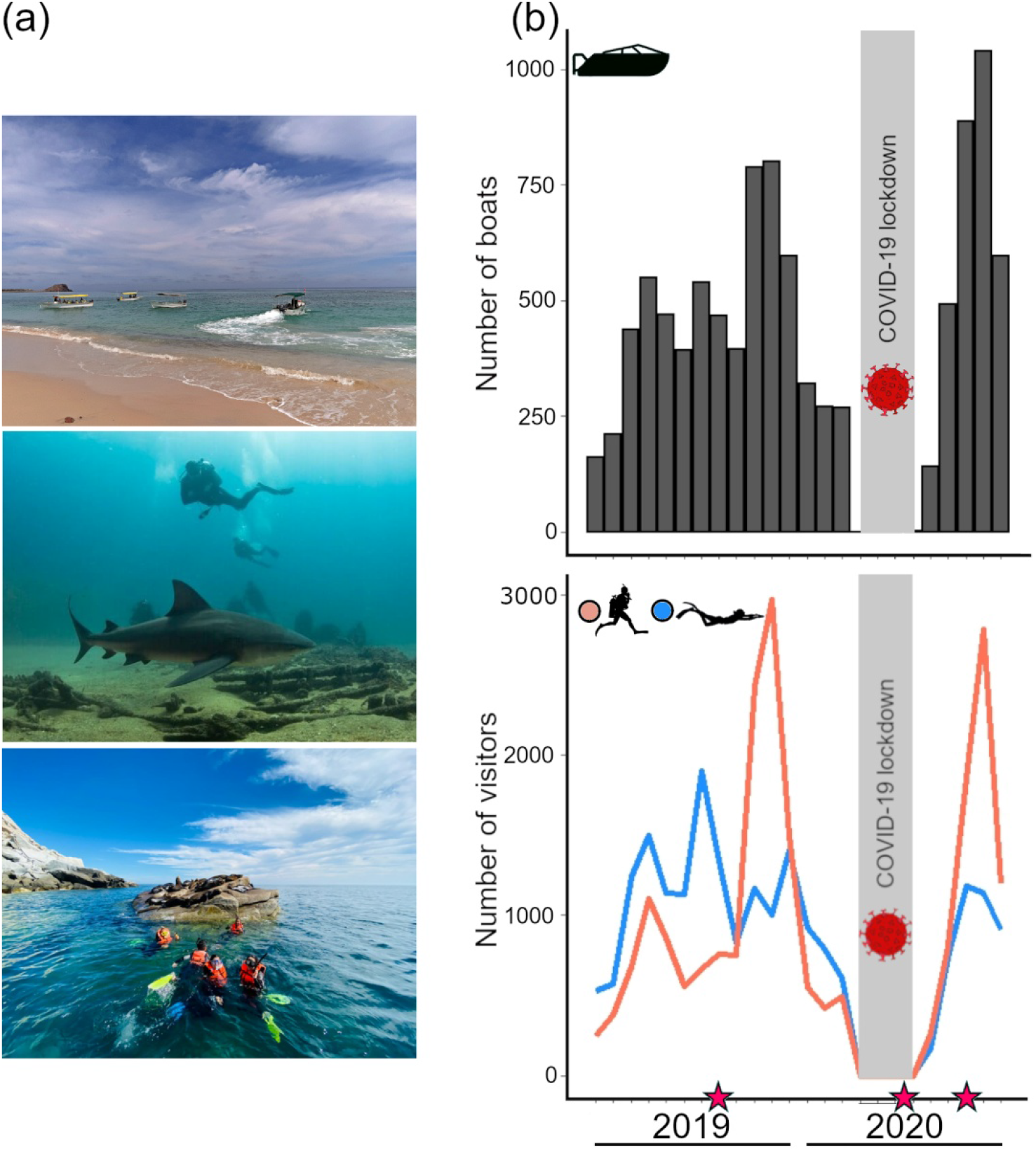
a) Ecotourism activities in Cabo Pulmo, photos by the authors and from the web (© Manta Scuba Diving). b) Number of boats, snorkelers, and divers entering the park monthly. Data from the “Comisión Nacional de Áreas Naturales Protegidas” (CONANP 2020).

During the first wave of the COVID-19 pandemic, the Mexican government closed all of the National Parks in the country. In Cabo Pulmo, only vessels serving ecotourism activities are allowed to enter the reserve. Therefore, the park’s closure from March 24 to August 15, 2020 reduced boat traffic inside the area to nearly zero (Figure 1b). We surveyed the reef fish assemblages in the marine reserve before, during, and immediately after prohibited public access. Thanks to this dataset, we found that the fish density more than doubled during the lockdown period. This knowledge may trigger further investigations on the disturbance(s) caused by apparent nature-friendly human activities on fishes’ behavior and lead to new management plans to improve the habitat quality of the marine reserves.

## Methods

### Data collection

We performed 145 underwater censuses using belt-transects of 100 m^2^ (4 x 25 m) at seven sites in Cabo Pulmo (Figure 2) in August 2019 (pre-lockdown period), June–July 2020 (lockdown period), and October 2020 (post-lockdown period). The Gulf of California has two distinct seasons, the cold season from the end of December to early June and the warm season from the end of June to early December. All of the surveys occurred during the warm season (June–July and August–October). Moreover, the reef fish assemblages form a seasonally stable community in the southwestern Gulf of California (Sánchez-Caballero *et al*. 2019). Six sites were selected to encompass the main reef area of the reserve, where touristic activity is high (Figure 2). One site was located at the reserve border and is not visited by tourists (Figure 2). The seven sites represent shallow and deep sites, i.e., three sites with a mean depth lower than 10 m and four sites with a mean depth between 10 and 20 m. We recorded the presence of every conspicuous fish along six to ten transects by site and period (see online data repository). For each transect, we reported the number of species (species richness), calculated the fish density (number of individuals per 100 m^2^), and estimated the evenness of the assemblages (the distribution of individuals by species) using the Pielou index (J). The evenness allows us to determine whether a change in fish density is not due to a small subset of species.

**Figure 2.**
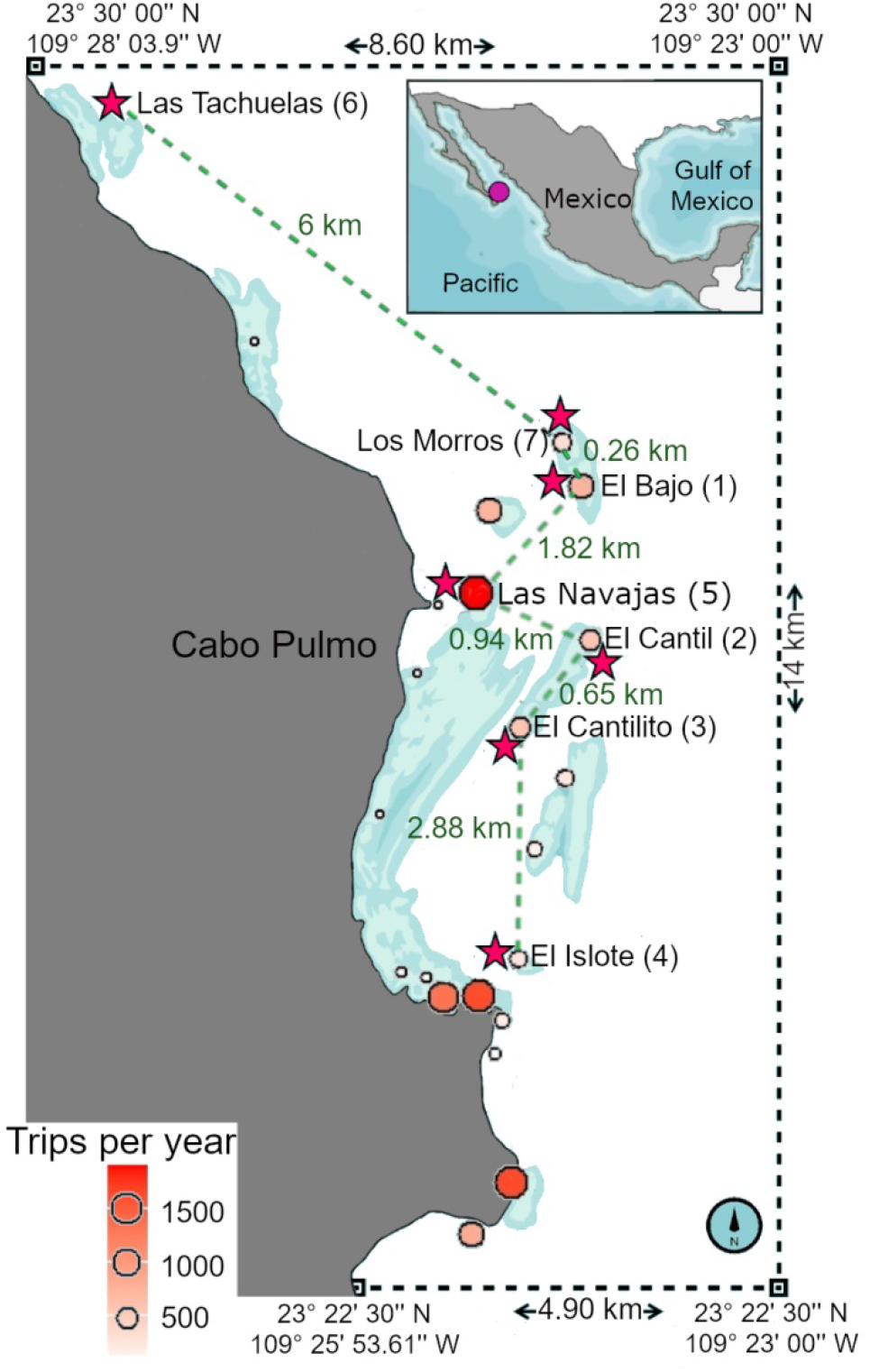
Map of Cabo Pulmo. Dashed lines represent the reserve boundaries; blue areas indicate the reefs; colored circles indicate sites visited by ecotourists. The size and color gradients of circles are proportional to the average number of visits by year (2017–2019); the pink stars represent the surveyed sites. Distances in kilometers between the surveyed sites are indicated.

### Statistical analysis

First, we ran linear mixed models (LMMs) to test the lockdown effect on the three diversity metrics of the fish assemblages (by transect). We log-transformed fish density to fit a normal distribution, and we used raw values for species richness and evenness. Sites and depth categories (from 3 to 10 m and >10 m) were added as random variables to avoid autocorrelation in the residuals. We did not detect violations of normality or variance homogeneity in the residuals of any of the three models.

Second, we determined which fish families changed in density during the lockdown, i.e., we compared the lockdown period to the pre-lockdown period. We focused on the most common families for statistical robustness, i.e., those observed in at least 20% of transects. For each family, we ran six models: generalized linear models (GLM) with a Poisson or negative binomial distribution, either with or without random variables (sites and depth category), and zero-inflated (ZI) models with a Poisson or negative binomial distribution. For each family, we selected the best of these models. We did not consider models with high residual overdispersion values (>1.5). When multiple models met the overdispersion condition, we selected the model with the lowest Akaike Information Criteria (we considered irrelevant differences lower than two and chose the simplest model in that case). Based on the selected models, we calculated the fitted fish density by family during and before the lockdown by bootstrapping with 1,000 iterations. We used the difference between the mean of the fitted abundances during each period and the standard error of this difference to calculate a t-value, as follows:

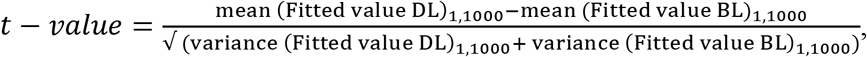

where DL and BL refer to “during” and “after” lockdown, respectively. Those t-values equal to or higher than two in absolute value indicated significant results at a 95% confidence interval (CI). We used a bootstrapping method to be able to compare the parameters (e.g., t-value) between distinct models (GLM(M) and ZI). We ran all analyses using R software (R Development Core Team 2016). The LMMs and GLM(M)s with negative binomial distribution were run using the R package *lme4* (Bates *et al*. 2018) and the ZI models using the R package *pscl* (Zeileis *et al*. 2008).

## Results

We observed a 2.5-fold increase in fish density in the absence of visitors against the period before the lockdown (411 [CI = 315–538] vs. 168 [CI = 124–229] fish by 100m^2^; Figure 3a, WebTable 1). To further support our results, we considered the historical monitoring data within the reserve. Two of the seven sites surveyed in the current study have been consistently monitored since 2007 during the warm season (El Cantil and Las Navajas). This long-term survey confirms the unusual fish density level observed during the lockdown period (WebFigure 1). The rise in fish density during the lockdown applied to the fish community overall, as the distribution of individuals among species (evenness) did not change during the lockdown (WebTable 1). Indeed, almost all of the 16 most common fish families (representing small, large, sedentary, and mobile fish, also belonging to different trophic groups) showed a clear trend of increasing density during the lockdown period (Figure 3b). The changes were significant in nine of them (the 95% CI of the estimate did not cross zero) (Figure 3b, WebTable 2).

**Figure 3.**
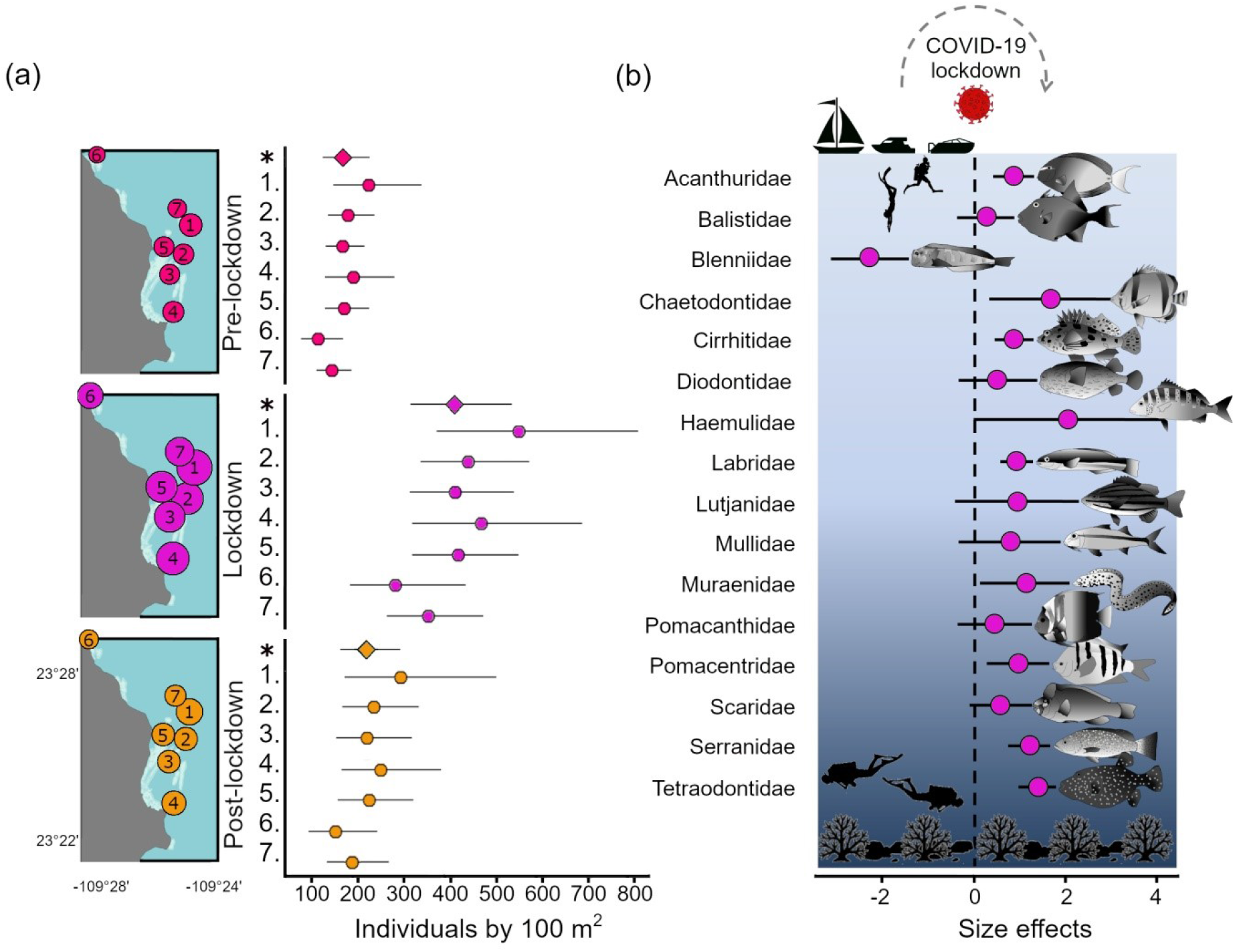
Increase in reef fish density during the COVID-19 lockdown. (a) Surveyed sites are plotted on the maps; circles sizes are proportional to fish density. The estimates of fish density (with 95% CI) are also shown for the three periods. Asterisks indicate the estimates for each period (considering sites and depth effects), and the numbers (1–7) indicate estimates for each site (see Figure 2 for full names). (b) Changes in fish density by family during the lockdown in comparison to the pre-lockdown period; the size effects (the difference between the two periods on a logarithmic scale) with 95% CI are illustrated.

The species richness also increased during the lockdown period (WebTable 1). It is noteworthy that during the lockdown period we observed some species rarely seen on the reef. Some of them are important in the aquarium trade, such as the endangered clarion angelfish (*Holacanthus clarionensis*) not seen in the area since 2005, the convict surgeonfish (*Acanthurus triostegus*), and the dragon wrasse (*Novaculichthys taeniorus*), as well as large commercial fishes like the jacks (Carangidae), longfin yellowtail and bluefin trevally (*Seriola rivoliana* and *Caranx melampygus*, respectively), and the broomtail and sawtail groupers (Serranidae; *Mycteroperca xenarcha* and *Mycteroperca prionura*,respectively).However, in the post-lockdown period, both species richness and fish density returned to the initial “normal” levels, although a slight trend of increased density was still observed (Figure 3a, WebTable 1).

## Discussion

During the lockdown period, the rise in fish density was as notable as its fall after the lockdown period. Once the reserve opened again, the touristic activities rapidly reached pre-lockdown levels (Figure 1b). Since the changes (rise and fall) happened in such a short time span, we can rule out larval recruitment or population displacement as causes of the shift in fish assemblages. Furthermore, we did not note unusual levels of juveniles during the lockdown period. We still know little about spawning aggregation in the Gulf of California (Sala *et al*. 2003), but we did not observe such behavior during the lockdown period. During the lockdown period, there was a worldwide concern about reserve managers, non-governmental organizations, and researchers’ capacity to access the reserves, increasing poaching risk (Corlett *et al*. 2020). However, Cabo Pulmo is a well-enforced marine protected area under strict control by the local community living inside the reserve. Thus, poaching is unlikely before, during, and after the lockdown. Moreover, a hypothetical increase in poaching during the lockdown would further support our findings, i.e., fish density would be even higher. Finally, a very recent study in French Polynesia also showed a more than 2-fold increase in fish density in ecotourism sites during the lockdown period of Bora-Bora, corroborating our findings (Lecchini *et al*. 2021). Therefore, we assert that ecotourism, the only authorized human activity in Cabo Pulmo, impacts the presence and behavior of marine fauna, even in a well-managed no-take reserve. Our findings agree with biomass and fish diversity estimation in remote wilderness areas, where human impacts are low (D’agata *et al*. 2016; McClanahan *et al*. 2019). Like remote zones, the fish diversity found during the lockdown may serve as a benchmark for management effectiveness (D’agata *et al*. 2016).

Pollution from oil, gasoline, and divers may drive noticeable and short-term changes in fish behavior (Armstrong *et al*. 2019; Samia *et al*. 2019). Boats used for tourism activities are well-maintained in Cabo Pulmo but scuba-diving may have a significant impact. Indeed, while the scuba-diving sites are much less visited than the most popular snorkeling spot, they showed similar or higher fish density increase during the lockdown (WebFigure 2). However, we have to investigate further the impact of these two activities on fish behavior. Nevertheless, the absence of divers or snorkelers in the water alone cannot explain the increase in fish density during the lockdown as the change was also observed in the only site not visited by tourists, even if it was in a slightly lower proportion than in ecotourist areas (WebFigure 2). Therefore, the reduced amount of noise from small vessels is likely a major factor explaining such changes in fish behavior. The sound in the water propagates rapidly in all directions and will affect all sites in the relatively small reserve, even those less-visited. The noise produced by small recreational and fishing boats or festive land-based human aggregations are known to affect fish behavior (Simpson *et al*.2016; Leduc *et al*. 2021), and the impact of human noise pollution (“anthrophony”) on marine wildlife is of great concern (Duarte *et al*. 2021).

Various families that increased in density during the lockdown period are sedentary or site-attached and cannot escape a noisy environment (e.g., Cirrhitidae, Diodontidae, Muraenidae, Pomacentridae). Under a usual noisy scenario (no lockdown), we hypothesize that a large proportion of fishes may remain hidden, and scuba diving censuses do not monitor them. The effects of noise pollution on fish behavior have only been quantified on a few reef fishes (Ferrier-Pagès *et al*. 2021). Nevertheless, a study on the juveniles of the damselfish *Pomacentrus wardi* showed that the species’ activity and boldness were half under the noise of 2-stroke outboards motor of that of fish under ambient reef noise conditions (McCormick *et al*. 2018). Noise pollution may also affect the survival rate of reef fishes, modifying population demographic (Simpson *et al*. 2016). Here, we don’t think that modification in the survival rate is a significant driver of the fish density increase observed during the lockdown due to the short period of time where such changes occurred. However, such disturbances should be monitored. The density of highly mobile and large-sized species also increased during the lockdown. It is difficult to believe that these hide during the scuba diving censuses (under a noisy scenario); instead, they may move to other more silent places. During the lockdown, access was prohibited to marine reserves and all of the beaches in the region. Therefore, all coastal reefs inside or outside the reserve should have been unusually quiet. For example, various testimonies revealed the presence of sharks along the coasts around Cabo Pulmo while they usually stay inside the reserve. Thus, the density increase of mobile species during the lockdown is likely not due to their displacement from “still noisy” adjacent reefs (outside Cabo Pulmo). Rather, we believe that mobile species could swim back and forth to adjacent deep zones to avoid disturbances from touristic activity and stayed, at least in larger proportion, on the reef during the lockdown. For example, *Lutjanus argentiventris, Mycteroperca rosacea*, and *Scarus ghobban* (the most common snapper, grouper, and parrotfish species in the region, respectively) are well-known to use the mesophotic zone (~40–50 m) (Sala *et al*. 2003; Cure *et al*. 2021).

Our results are of great importance for marine conservation and management because they show that ecotourism has direct consequences on the behavior of ichthyofauna. The ecotourism sector, essential for the functioning of the marine reserves, needs to be adequately regulated to preserve wildlife and the decent economic income it offers to local populations (Sala and Giakoumi 2018). Nevertheless, our results also show that fish density in no-take reserves with high human activity may be much higher than currently estimated. This observation provides a great incentive to further improve the health of protected areas by addressing the problem of ecotourism impact. For example, little is currently being done to tackle noise pollution in the marine environment (Markus and Sánchez 2018). The COVID-19 lockdown revealed that small motorboats’ noise pollution might affect fish movement and habitat use and highlighted the need to monitor anthropogenic noise in reserves’ management plans. We cannot yet quantify from which noise level a large panel of species may be affected but such knowledge is essential to improve marine reserve management. Indeed, noise pollution would not only induce underestimation of fish diversity but may have various negative effects on fish health and ecosystem functioning by affecting fish distributions, fitness, species interactions, and masking sound communication of numerous vocal species (Slabbekoorn *et al*. 2010). Therefore, we encourage future research and managers’ involvement to determine the threshold or the type of anthropogenic sound affecting the behavior of fishes.

We provide some perspectives to further investigate the effects of sound pollution on fish assemblages in marine reserves. The sensitivity to noise disturbance will vary among species (Popper and Hawkins 2019). Therefore, behavioral studies following the activity of several species *in situ* while recording the ambient noise could shed light on the effect of motorboat sound disturbance, similarly to some previous studies made on damselfishes (Simpson *et al*. 2016; McCormick *et al*. 2018). For mobile species, acoustic telemetry technology can be used to correlate the movement of the fishes with the levels of motorboats’ noise (Barcelo-Serra *et al*. 2021). Although high-resolution telemetry represents an efficient tool to study fish responses to environmental perturbations (Aspillaga *et al*. 2021), it can be expensive, difficult to put in place, and require the precise monitoring of numerous individuals (Barcelo-Serra *et al*. 2021). On the other hand, many reef fish species are well-known to be vocal, including various mobile fish families, e.g., Haemulidae, Lutjanidae, Scaridae, and Serranidae (Bertucci *et al*. 2014; Parmentier *et al*.2021; Tricas and Boyle 2021). The sounds produced by the fish and other organisms (“biophony”) can be used to track their activity (Tricas and Boyle 2021). Studies relating biophony with anthropogenic sound, called soundscape ecology (Pijanowski *et al*. 2011), can be a powerful, cost-effective tool for reserve management (Ferrier-Pagès *et al*. 2021). Network of hydrophones could be positioned over extended periods, inside and outside the reserve, at different depth ranges, and considering a range of use rate (site popularity) to record the activity of fishes and determine a relationship with anthropogenic sound pressure.

To conclude, better knowledge about noise pollution on fish behavior can lead to direct reserve managing actions.. On one hand, the reserve could be temporarily closed but at the risk of encountering strong opposition by the local population managing the reserve unless financial compensations is offered, which is unlikely in emergent and developing countries. So, such drastic solutions may be counter-productive as human population adherence is essential. To reduce the number of boats could also be an unpopular decision. On the other hand, some regulations as simple as speed reductions can be settled (Mccloskey *et al*. 2020). Technological solutions also exist for a quieter marine environment, e.g., the use of 4-stroke instead of 2-stroke engines affected the behavior of a damselfish species drastically less (McCormick *et al*. 2018), and electric engines or specifically designed propellers can be much less noisy (Jones 2019; Parsons *et al*. 2020). Contrary to global change problems, noise pollution is a temporal disturbance that we can turn off. A whole range of tools already exists to have quieter marine reserves and provide higher quality habitats to the community that they host.

## Acknowledgments

Funding to conduct field work was provided by the Universidad Autónoma de Baja California Sur (UABCS) and the following civil associations: Costa salvaje A.C., Sociedad de Historia Natural Niparajá A.C., Amigos para la Conservación de Cabo Pulmo, Baja Coastal Institute and Pronatura Noroeste. We thank Romeo Saldívar-Lucio, Juan Carlos Perusquía-Ardón (Centro de Investigación Científica y de Educación Superior de Ensenada), Arturo Ayala-Bocos (Ecosistemas y Conservación, A.C.) and local dive guides, who collaborated in the fish census. We thank Luis Mario Castro for sharing photos. Permission to perform the study was granted by Parque Nacional Cabo Pulmo (Carlos Godínez-Reyes, Director of the park). We thank the people from Cabo Pulmo, who were graceful enough to allow us to visit the town in the midst of the pandemic event. We thank L Alvarez-Filip, B. Frédérich, and L. Kever for their helpful discussions and comments on the manuscript.

---

**WebFigure 1.**
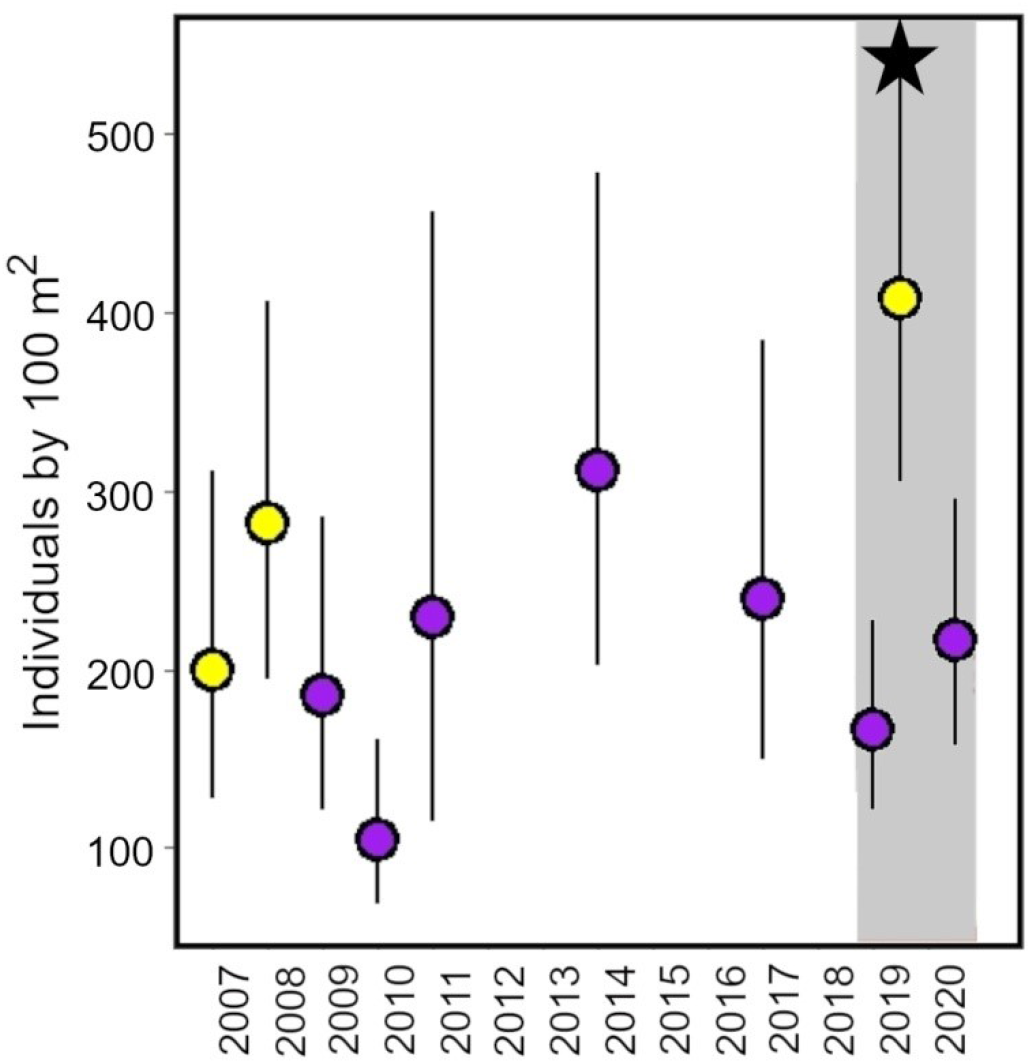
Estimated fish density (+/- standard error) in Cabo Pulmo from 2007 to 2020. Linear mixed models (considering the sites as a random variable) estimated the fish densities thanks to the data of two different sites (El Cantil and Las Navajas) consistently monitored since 2007. All data were collected during the warm season in the Gulf of California (end of June to December). The yellow and purple circles represent the periods June–July and August–October, respectively. The grey rectangle represents the period studied in the main manuscript, i.e., before, during, and after the lockdown. The black star represents the fish density estimated during the lockdown period. Although monitoring is usually not performed during the June–July period, it does not seem to present higher fish density than the August–October (except when the lockdown of the reserve occurred).

**WebFigure 2.**
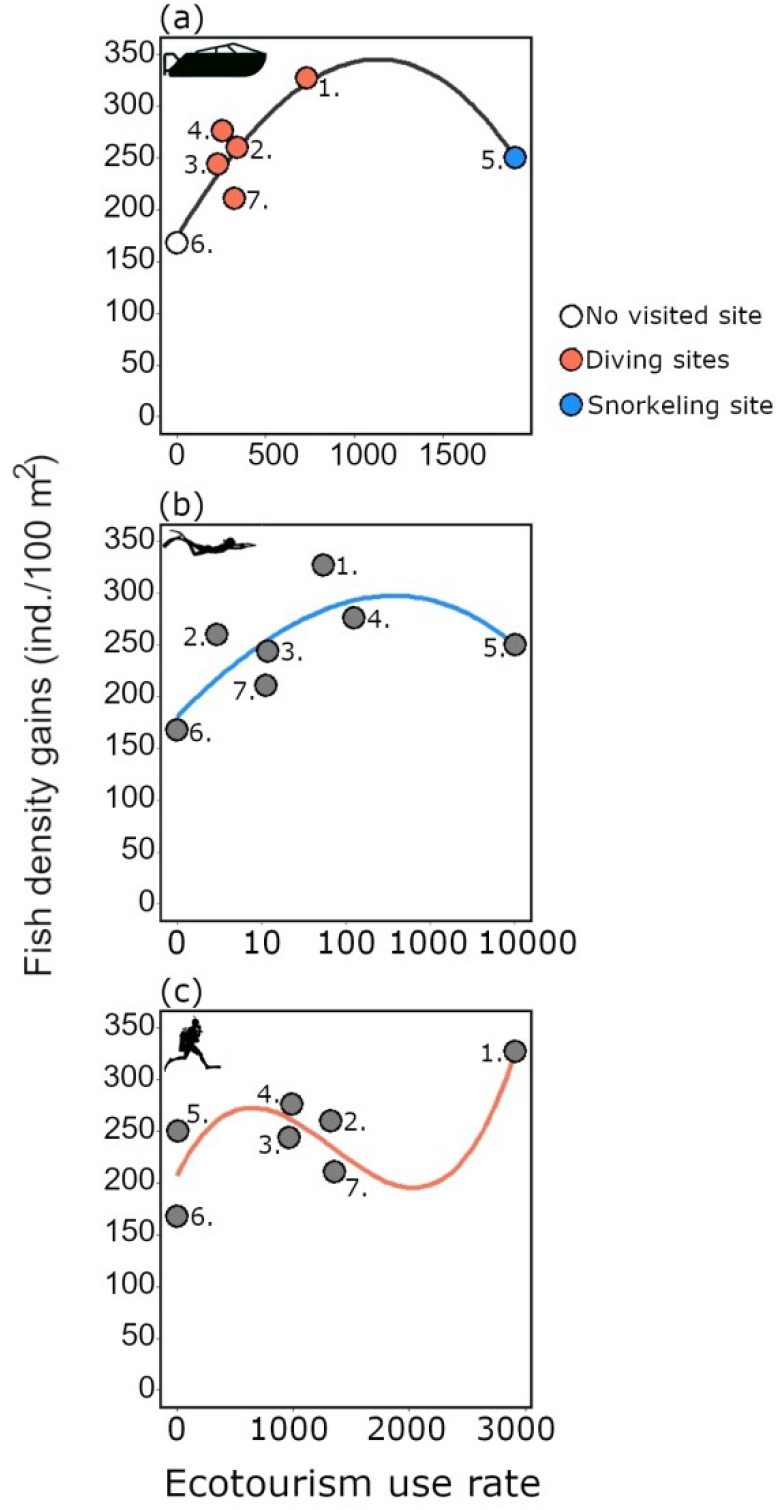
The relationship between the gains in fish density (difference between during and before the lockdown) and a) the number of boat trips, b) snorkelers (log scale), and c) divers per year (averaged for the period of 2017–2019) in the seven surveyed sites. The lines are the result of third-order polynomial regression. Each site is numbered as on the map of Cabo Pulmo (Figure 2).

**WebTable 1.** Comparison of the three fish diversity metrics during and after the lockdown to before the lockdown. Linear mixed models with the sites and the depth as random variables were used. The estimate (difference between during/after the lockdown to before the lockdown), the standard error (SE), the t-value, and the random standard deviance (RSD) of the two random variables are shown

|  |  | Estimate | SE | t-value | RSD<br>Site | RSD<br>Depth |
| --- | --- | --- | --- | --- | --- | --- |
| Species richness |  |  |  |  | 1.08 | 0.83 |
|  | Before lockdown (Intercept) | 14.61 | 0.95 | 15.33 |  |  |
|  | During lockdown | 2.83 | 0.81 | 3.48 |  |  |
|  | After lockdown | -0.40 | 0.89 | -0.45 |  |  |
| Abundance (log) |  |  |  |  | 0.23 | 0 |
|  | Before lockdown (Intercept) | 5.12 | 0.15 | 33.47 |  |  |
|  | During lockdown | 0.90 | 0.16 | 5.51 |  |  |
|  | After lockdown | 0.26 | 0.18 | 1.48 |  |  |
| Evenness |  |  |  |  | 0 | 0 |
|  | I Before lockdown (Intercept) |  |  |  |  |  |
|  | During lockdown | -0.04 | 0.03 | -1.36 |  |  |
|  | After lockdown | -0.09 | 0.03 | -2.62 |  |  |

**WebTable 2.**
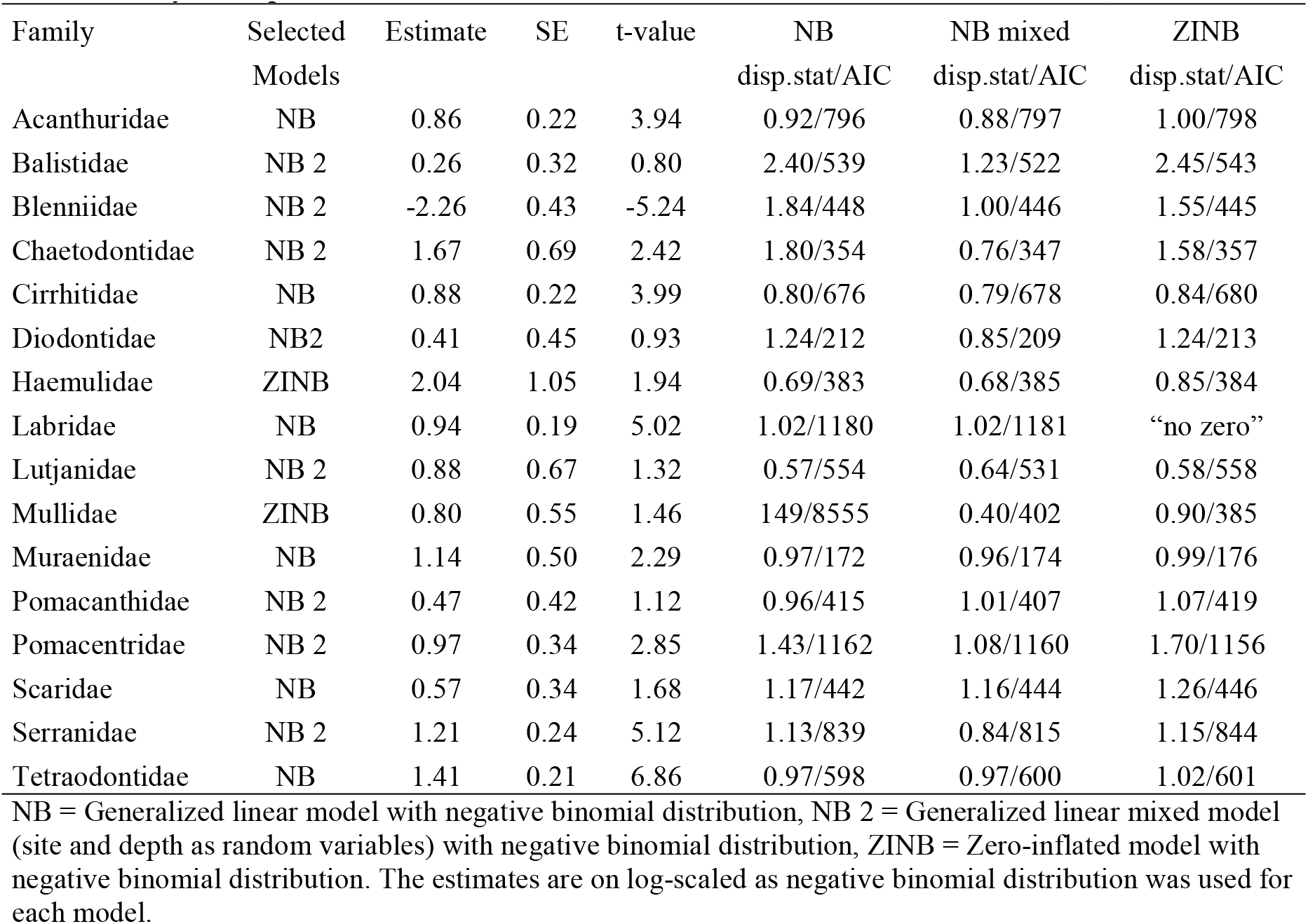
Outputs of the models run with 1000 bootstraps to compare the fish density by family between before and during the lockdown. The estimate (difference between the two groups), the standard error (SE) and the t-value are shown. For each family, residuals overdispersion (disp.stat) and Akaike (AIC) values were indicated for the different tested models. Only models selected at least for one family are represented.

| Family | Selected Models | Estimate | SE | t-value | NB<br>disp.stat/AIC | NB mixed<br>disp.stat/AIC | ZINB<br>disp.stat/AIC |
| --- | --- | --- | --- | --- | --- | --- | --- |
| Acanthuridae | NB | 0.86 | 0.22 | 3.94 | 0.92/796 | 0.88/797 | 1.00/798 |
| Balistidae | NB 2 | 0.26 | 0.32 | 0.80 | 2.40/539 | 1.23/522 | 2.45/543 |
| Blenniidae | NB 2 | -2.26 | 0.43 | -5.24 | 1.84/448 | 1.00/446 | 1.55/445 |
| Chaetodontidae | NB 2 | 1.67 | 0.69 | 2.42 | 1.80/354 | 0.76/347 | 1.58/357 |
| Cirrhitidae | NB | 0.88 | 0.22 | 3.99 | 0.80/676 | 0.79/678 | 0.84/680 |
| Diodontidae | NB2 | 0.41 | 0.45 | 0.93 | 1.24/212 | 0.85/209 | 1.24/213 |
| Haemulidae | ZINB | 2.04 | 1.05 | 1.94 | 0.69/383 | 0.68/385 | 0.85/384 |
| Labridae | NB | 0.94 | 0.19 | 5.02 | 1.02/1180 | 1.02/1181 | “no zero” |
| Lutjanidae | NB 2 | 0.88 | 0.67 | 1.32 | 0.57/554 | 0.64/531 | 0.58/558 |
| Mullidae | ZINB | 0.80 | 0.55 | 1.46 | 149/8555 | 0.40/402 | 0.90/385 |
| Muraenidae | NB | 1.14 | 0.50 | 2.29 | 0.97/172 | 0.96/174 | 0.99/176 |
| Pomacanthidae | NB 2 | 0.47 | 0.42 | 1.12 | 0.96/415 | 1.01/407 | 1.07/419 |
| Pomacentridae | NB 2 | 0.97 | 0.34 | 2.85 | 1.43/1162 | 1.08/1160 | 1.70/1156 |
| Scaridae | NB | 0.57 | 0.34 | 1.68 | 1.17/442 | 1.16/444 | 1.26/446 |
| Serranidae | NB 2 | 1.21 | 0.24 | 5.12 | 1.13/839 | 0.84/815 | 1.15/844 |
| Tetraodontidae | NB | 1.41 | 0.21 | 6.86 | 0.97/598 | 0.97/600 | 1.02/601 |

NB = Generalized linear model with negative binomial distribution, NB 2 = Generalized linear mixed model (site and depth as random variables) with negative binomial distribution, ZINB = Zero-inflated model with negative binomial distribution. The estimates are on log-scaled as negative binomial distribution was used for each model.

